# PICS: Pathway Informed Classification System for cancer analysis using gene expression data

**DOI:** 10.1101/047514

**Authors:** Michael Young, David Craft

**Affiliations:** Department of Radiation Oncology, Massachusetts General Hospital, Harvard Medical School

## Abstract

We introduce PICS (Pathway Informed Classification System) for classifying cancers based on tumor sample gene expression levels. PICS is a computational method capable of expeditiously elucidating both known and novel biological pathway involvement specific to various cancers, and uses that learned pathway information to separate patients into distinct classes. The method clearly separates a pan-cancer dataset into their tissue of origin and is also able to sub-classify individual cancer datasets into distinct survival classes. Gene expression values are collapsed into pathway scores that reveal which biological activities are most useful for clustering cancer cohorts into sub-types. Variants of the method allow it to be used on datasets that do and do not contain non-cancerous samples. Activity levels of all types of pathways, broadly grouped into metabolic, cellular processes and signaling, and immune system, are useful for separating the pan-cancer cohort. In the clustering of specific cancer types, certain pathway types become more valuable depending on the site being studied. For lung cancer, signaling pathways dominate, for pancreatic cancer signaling and metabolic pathways, and for melanoma immune system pathways are the most useful. This work suggests the utility of pathway level genomic analysis and points in the direction of using pathway classification for predicting the efficacy and side effects of drugs and radiation.

## 1 Background

Cancer is a genetically heterogeneous disease, both across cancer types and within. Nevertheless, most patients are prescribed treatments without regard to any specific biological signatures of their disease. Exceptions to this generally take the form of drug selection based on a single genetic mutation (examples include venurafenib for BRAFv600 mutations, erlotinib for EGFR mutations, and crizotinib for ALK mutations) or in limited research settings based on a signature derived from a small number of genes (anywhere from one to about 50; see e.g. [1, 2]). Such genomic-based partitioning of patients fails to predict with appropriate levels of confidence whether a potentially highly toxic drug will be effective or what side effects, and their intensities, may occur. This issue is compounded by the fact that advanced therapies are designed to target specific pathways, meaning treatments dependent on pathway regulation are being prescribed with incomplete knowledge of the state of the pathway itself.

A typical scenario for developing a predictive model for cancer treatment involves a small cohort of patients, usually less than 100, and gene expression data for each of them with upwards of 20,000 gene expression levels measured. In this setting, one must be careful to avoid correlations that appear due to chance alone [3]. One way to deal with this is to collapse gene-level data into more compact, functional pathway-level data. A biological pathway is a set of biochemical reactions that perform a specific nameable function. A classic example of a metabolic pathway is the production of ATP from glucose. Pathways have been discovered through decades of laboratory research and are systematically curated in several publicly accessible resources, including KEGG [4], BioCarta [5], Reactome [6], and WikiPathways [7]. Several research teams have developed techniques to use pathways to aid in the interpretation of gene expression data. Some of the methods output a score indicating how dysregulated each pathway is for each patient (e.g. [8, 9, 10, 11, 12, 13, 14]). Of these Pathifier [10] and Kim-DeLisi [14] additionally use the pathway scores to classify patients. However, these algorithms are encumbered by computational challenges and limited scope, and thus have not yet been successfully applied to actually improve treatment response prediction.

We developed the Pathway-Informed Classification System (PICS) to extend the utility of pathway-based approaches to cancer classification with a method that is fast, robust, and scalable to large patient cohorts across all available cancer types. We validate the method on both pan-cancer cohorts, where we show separation of cancer types, and on individual cancer type cohorts, where we show that the patient clusters found split the patients into statistically distinct survival classes. This work serves as a stepping stone to a biologically-informed statistical approach to predicting patient response to drugs and radiation therapy.

## 2 Methods

PICS (Pathway Informed Classification System) consists of two steps: the first step assigns to each patient and each biological pathway a score which is either a single scalar value or a small set of numbers, and the second step takes these pathway scores for the entire cohort of patients and uses them to cluster the patients.

The input to PICS is a matrix of messenger RNA (mRNA) expression values for a large number of genes for all the patients of a cohort. We require the matrix to be full and that all the patient samples, which may include patients from across disease types and non-cancerous matched tissue samples as well, come from the same microarray technology so that the values are comparable. We will discuss the incorporation of other types of genomic data as well but for this report we use only mRNA expression level data, which we obtain from publicly available resources.

We obtain pathway descriptions from the KEGG database API, which is also publicly available. The KEGG pathway database contains 301 pathways related to humans. The definition of a biological pathway, in particular the boundaries of the pathways, are not well agreed on, and indeed some KEGG pathways are combinations of more primitive KEGG pathways. Some pathways have been attributed to a specific cancer (e.g. KEGG HSA:05216, thyroid cancer pathway) when really those pathways contain gene interactions that are implicated in cancer in general. The set of pathways used will have an effect on the results of the PICS algorithm, but as a first pass we start by including all possible pathways from a given resource (KEGG in our case) without regard to the pathway’s speculated involvement in cancer. We also use KEGG modules, which are descriptions of more basic reactions and therefore generally involve fewer genes. The KEGG database contains 187 modules for humans.

### 2.1 PICS step 1: scoring pathways

For our purposes, a pathway or module will be represented by the set of genes in that pathway. We ignore the network topological structure of the pathway and instead view the pathway as a set of genes. Since our goal is an an improved classification and predictive framework, we believe this representation is an appropriate level of detail, where biology in terms of closely interacting gene products is taken into consideration but detailed interactions are ignored [10]. It would not be hard to include the network structure of the pathways however, as has been done [8, 9, 11]. The only requirement for our pathway scoring is to take the gene expression data for pathway genes and condense this information into a single number or small set of numbers. For this we introduce three methods: PCA, NTC, and GED.

#### 2.1.1 PCA – principal components analysis

The simplest way to reduce the dimensionality of a set of gene expression levels across a set of patients is with PCA[15]. Using PCA also provides a straightforward means to condense the expression data to either a scalar (i.e. the first principal component, that is, the projection of each patient’s gene expression values onto the vector representing the first principal component) or a vector which consists of the first s principal component projections. The number of PCs used in a specific pathway scoring is user-configurable. The user can input either a fixed integer value or a fraction *f*, which gets interpreted as *round*(*f* × *I_D_*) where *I_D_* is the intrinsic dimensionality estimate [16] of the pathway data matrix.

The PCA method makes no assumptions about the underlying datasets and is thus applicable to pan-cancer cohorts, datasets that include normal tissue samples, and single cancer type datasets that contain just cancer samples. For sets that do have normal tissue samples, we provide two specialized methods, NTC and GED, which score pathways based on the differences of their gene expression values compared to the normal samples.

#### 2.1.2 NTC – normal tissue centroid

For a set of *g* genes (the genes of a particular pathway) each tissue sample can be viewed as a point in *g*-dimensional space. The NTC, or normal tissue centroid, is the location in this space computed by averaging the coordinates of each of the normal tissues. One can then score each sample by computing its distance in *g*-space from the NTC. This method is similar to the Pathifier method which, for each sample, computes the distance away from the normal samples along a curved line passing through the normal samples [10].

#### 2.1.3 GED – gene expression deviation

The PCA and the NTC methods do not model the fact that some genes are over-expressed and some are under-expressed in the cancerous samples, in a patient-specific manner. The GED, or gene expression deviation, score gives two numbers for each patient and each pathway: an aggregate over-expression score and an aggregate under-expression score. For each pathway, pathway genes are selected for inclusion in this score based on how the distribution of expression values for the cancer samples differs from those of the normal samples. Specifically, the Kolmogorov-Smirnov test for the difference of distributions is used. If a gene *g* passes the test, meaning its gene expression values are distributed significantly differently from the normal to the cancerous tissues, then for each patient *p* and that gene *g*, we form the differential-expression score Δ*_pg_* as:

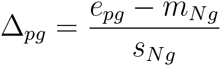

where *e_pg_* is the expression level for patient *p* of gene *g, m_Ng_* is the mean expression level of gene *g* for the normal samples and *s_Ng_* is the standard deviation of the expression levels across the normal samples. Then for each sample we form two scores: one which adds up the positive values of Δ*_pg_* across the pathway genes, and one which adds up the negative values.

#### 2.1.4 Deciding which genes to include

Many genes are involved in multiple pathways. For example, MAPK, PI3K, and AKT genes are all involved in over 60 KEGG pathways. Therefore the gene expression levels of these genes may not be closely tied to the activity of any one of its particular pathways. On the other hand, most pathways contain at least a few genes that are unique to that pathway. For all scoring methods, we investigate which gene expressions should go into a pathway score based on how many other pathways the gene belongs to. In particular, we define a parameter called Mcut, for membership cutoff: a gene is only included in a pathway score if that gene belongs to Mcut or fewer pathways.

#### 2.1.5 Deciding which pathways to include

Depending on the scoring technique, we can use various methods to decide whether or not to include a particular pathway in the matrix used for clustering.

For datasets containing normal tissue samples, we implement the PCA compression of the pathway as the first step in the selection process. Using the PCA representation we compute silhouette scores for each sample [17] by using the groups normal and cancerous. For each sample the silhouette score is a measure of how well that sample fits into its defined group, given the data used for the clustering, which is the PCA compression. The maximum possible silhouette score for a sample point is 1, meaning the sample perfectly belongs to the rest of the samples in that group. The average of all the silhouette scores gives a measure of how well the normal and cancerous samples cluster. Only pathways that have an average silhouette score above a given threshold are included in the final pathway score matrix.

For datasets that do not have normal samples we rely on the patient cohort survival data to judge a pathway’s usefulness. For these datasets the PCA compressed pathway is clustered into two groups using either k-means or k-medoids clustering [18]. Given these two clusters, a pathway is accepted if the Kaplan-Meier survival curves for these groups separate at a p-value that is lower than a user-defined threshold.

### 2.2 PICS step 2: patient classification based on pathway scores

The above pathway selection and scoring results in the conversion of the gene expression-patient sample matrix, which we call m (*m* is of size [number of genes] × [number of patients]) into a pathway score matrix which we call *p*. If each pathway is represented for example by two principal components, and *h* is the number of pathways that pass the silhouette score, then the size of the matrix *p* will be [2*h*] × [number of patients]. Note that in general 2*h* << [number of genes], which means we have regularized or compressed the gene expression data.

We use three types of clustering to form distinct patient groups: k-means, k-medoids, and hierarchical. K-means and k-medoids are useful for fixing the number of clusters, for example two, which makes for easier interpretability of separation of Kaplan-Meier curves. K-medoids, where the cluster centers are chosen as the the “median” sample point of the cluster rather than the weighted average, i.e. mean location, is more stable against outliers in the data. Hierarchical clustering is a more natural grouping of patients that does not involve pre-specifying the number of groups. For hierarchical clustering we use UPGMA (Unweighted Pair Group Method with Arithmetic mean) implemented in the Matlab (Version R2015b, Natick MA) function clustergram.

### 2.3 Data sources

The pan-cancer set contains six datasets consisting of 568 patients for five cancer types (acute myeloid leukemia, kidney, adult germ cell, osteosarcoma, and ovarian) obtained from the PRE-COG database (Prediction of Clinical Outcomes from Genomic Profiles, [19]). Two sets are from different ovarian studies, to verify that ‘like’ cancer types from different studies cluster together using our method. All sets were profiled used the same microarray, the Affymetrix U133A, to minimize cross-set variability. We use all 301 pathways from KEGG for this dataset and PCA scoring with 2 PCs per pathway. The adult germ cell dataset includes six normal tissue samples. None of the other sets include normal samples.

We also downloaded and normalized six datasets of cancers with normal tissue from the NCBI database via PRECOG: localized pancreatic duct adenocarcinoma[20] (GSE21501), non-small cell lung carcinoma[21] (GSE19188), adult germ cell carcinoma and seminoma[22] (GSE3218), ovarian[23] (GSE26712), glioblastoma[24] (GSE13041), and sub-optimally debulked ovarian (GSE-26712). The adult germ cell, ovarian, glioblastoma, and sub-optimally debulked ovarian datasets were profiled with the Affymetrix Human Genome U133A array. The pancreas set was profiled using the Agilent-014850 Whole Genome microarray. The lung set was profiled with the Affymetrix U133Plus array. Datasets varied in number of normal tissue samples, number of cancerous tissue samples, and number of genes reported, see Table 1. We also include one dataset which does not have normal samples present{a melanoma cohort of 470 patient samples. This is a TCGA dataset obtained via the Broad Institute’s GDAC firehose (gdac.broadinstitute.org, Skin Cutaneous Melanoma).

**Table 1:** Information for the individual disease site classifications.

| Cancer type | Source | Dataset | Genes in expression matrix | Number of Normal sample | Number of cancer samples |
| --- | --- | --- | --- | --- | --- |
| Pancreatic ductal adenocarcinoma | PRECOG | GSE21501 | 20936 | 30 | 102 |
| Non-small cell lung carcinoma | PRECOG | GSE19188 | 42423 | 65 | 82 |
| Adult germ cell carcinoma and seminoma | PRECOG | GSE3218 | 20936 | 6 | 74 |
| Ovarian | PRECOG | GSE26712 | 20965 | 10 | 185 |
| Glioblastoma | PRECOG | GSE13041 | 42391 | 4 | 80 |
| Sub-optimally debulked ovarian | PRECOG | GSE26712 | 20965 | 10 | 95 |
| Melanoma | TCGA | NA | 20512 | 0 | 470 |

For the six cohorts with normal samples present we loop through various scoring methods (PCA, NTC, GED), clustering types (k-means, k-medoids), pathway silhouette cutoffs, gene-pathway membership cutoff Mcut, and the number of principal components used, in order to find the parameter set that best clusters the patients. After a coarse parameter search we fine tune the parameters within that range to approach the `optimal’ solution, which is determined by Kaplan-Meier curve separation via two-group clustering. The result is a reduced, cancer-specific pathway score matrix which provides functional biological information and improved survival clustering.

We analyze a melanoma dataset from TCGA consisting of 470 samples across disease stages. Since this patient cohort was large, and since the pathway selection criteria, due to the lack of normal samples, was done based on the ability of the pathway to separate the KM curves, we split this dataset into a training and test set in order to gain more confidence in the robustness of the resulting classification. Similar to the classification of the individual cancer type datasets that had normal tissues, we loop through a wide range of parameter settings to find the best set of parameters to cluster the melanoma dataset. We additionally repeat the run for each parameter set forty times, each time randomly splitting the cohort into a training set and a test set, and we finally choose the parameter set that consistently, across most of the forty runs, effectively (regarding KM separation) clusters patients in both sets. We searched over the respective sizes of the training and test sets and found that a 50-50 split was optimal (this splitting was done for each of the runs by flipping a weighted coin for each patient, so the training and test group sizes are not always the same across the runs). Since there is a trade-off between how well you can separate the training set and how well you generalize to the testing set, we chose a point on the Pareto trade-off curve between these two objectives that balanced the two goals on average.

The optimal parameter set found was NumPCs=2 and the cutoff value for the p-value of the KM curve separation for each pathway of 0.05. For displaying the results, we choose the median result (regarding the p-value for the Kaplan-Meier separation of the training set) over the forty replicates.

## 3 Results

### 3.1 Pan cancer clustering

Figure 1 demonstrates the successful hierarchical clustering by disease type. Pathways generally sort by type which therefore allows us to add broad labels to the pathway groupings. See the Supplementary Information for the names of the pathways, from the top row downward. For example, all but one metabolic pathway is contained within the first grouping. This initial check validates that PCA can sort cancer types using pathway information via gene expression data. The two ovarian sets clustered together, and the normal tissues for the AGC normal samples clustered alongside the cancerous AGC samples (they are the six leftmost columns in that block). The most similar two cancer types from this viewpoint are kidney and AGC. AML and ovarian are similar regarding metabolic and cell processes pathways but differ in their immune and endocrine signature. Since PCA decomposition was used to score these pathways, the absolute values of the pathway scores are not biologically meaningful (principal components are arbitrarily oriented; the magnitudes of the values are relevant but not the signs). In order to assess the differences across cancers regarding a particular pathway, we can plot the gene expression levels for the genes in that pathway across the five cancer types. As an example, gene expression levels for the B-cell receptor signaling pathway are shown in Figure 2. This confirms the data in the pan cancer clustergram by showing that for this immune pathway, AML and kidney are similar, osteosarcoma and ovarian are similar, and AGC falls in between. Additionally, Figure 2 reveals that AML and kidney have the highest gene expression levels for this pathway overall and osteosarcoma and ovarian have the lowest.

**Figure 1:**
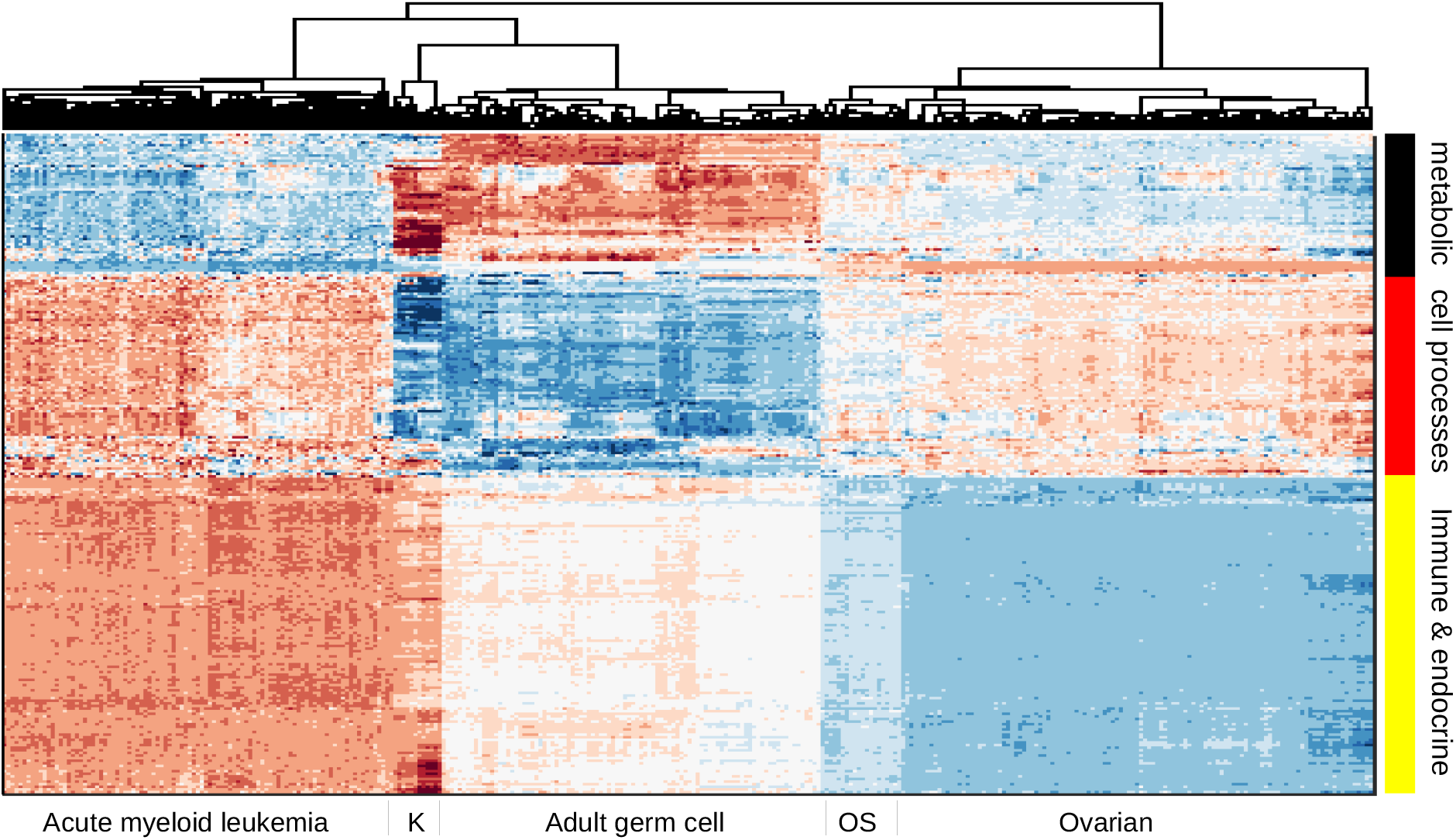
Hierarchical clustering across all pathways (scored by PCA method with two PCs per pathway) of five cancer cohorts combined into a single gene expression matrix. This figure serves as a proof of concept of pathway-based clustering by demonstrating the successful grouping of patients into their respective disease types via their gene expression levels. The hierarchical clustering results are shown in the upper tree dendrogram diagram, which has been suppressed for the pathway (row) sorting. K=kidney and OS=osteosarcoma. The pathway list for this heatmap appears in Supplementary Information.

**Figure 2:**
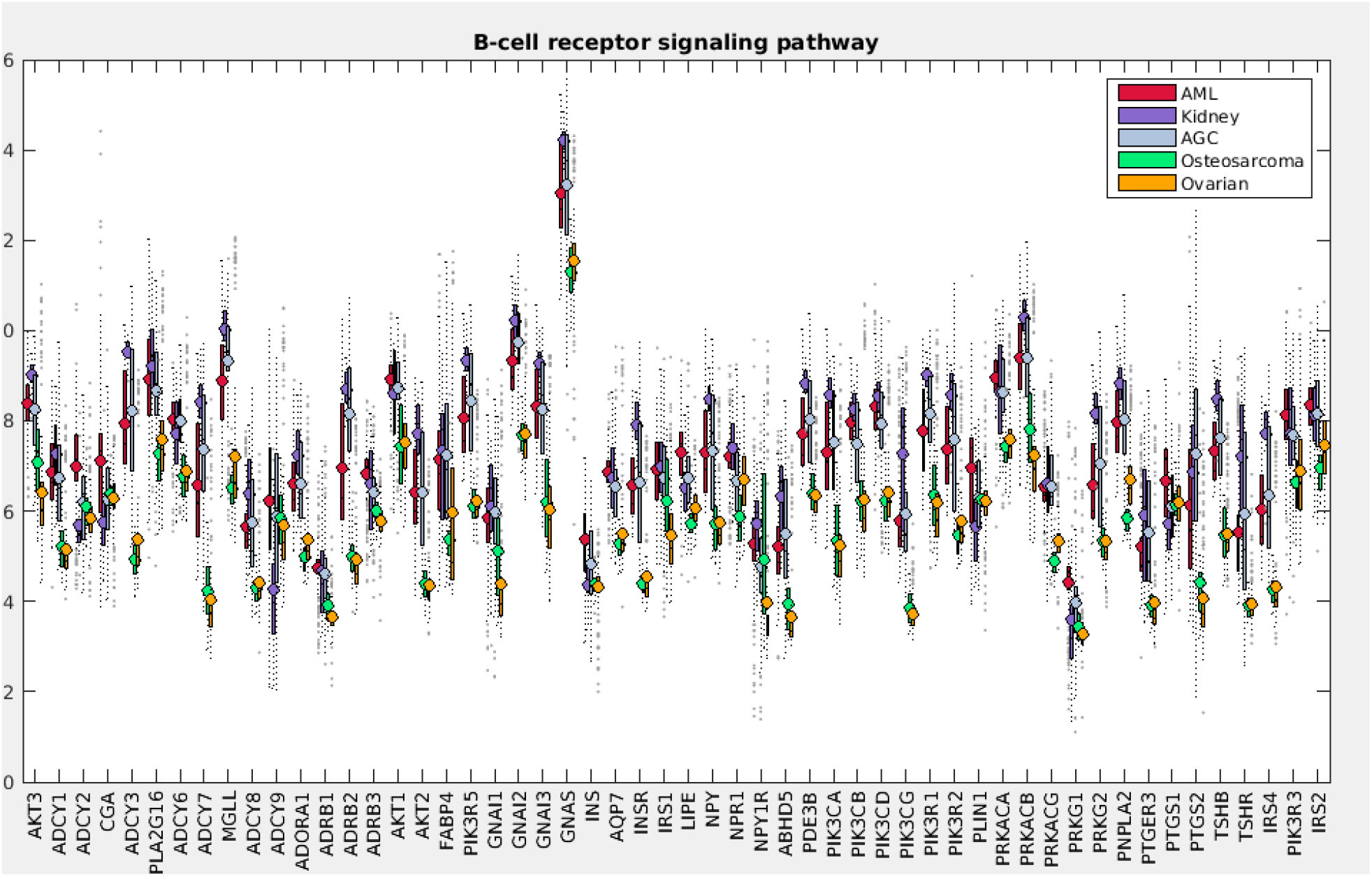
B-cell receptor signaling pathway gene expression levels for pan cancer analysis. Levels are directly from the pan cancer gene expression datasets, which have been log2 normalized.

### 3.2 Individual cancer site clustering with normal tissue present

Table 2 summarizes the runs and displays the best parameter sets/scoring types for each case. All six patient cohorts reached statistically significant solutions with p-values < .05. Pancreas and GBM were the most difficult to separate according to their p-values of 0.03. The sub-optimally debulked ovarian set had the best separation, per its low p-value (.0008). This is an improvement over an earlier gene expression-oriented study of the same dataset, which had a p-value of .02 [25] (the other studies did not produce patient groupings with Kaplan-Meier separation so no comparisons can be made). There is a general trend towards low p-value (good separability) and low number of pathways and modules used in the classification, however the pancreas dataset defies the trend being both relatively hard to separate but using very few pathways for optimal separation (discussed below).

**Table 2:**
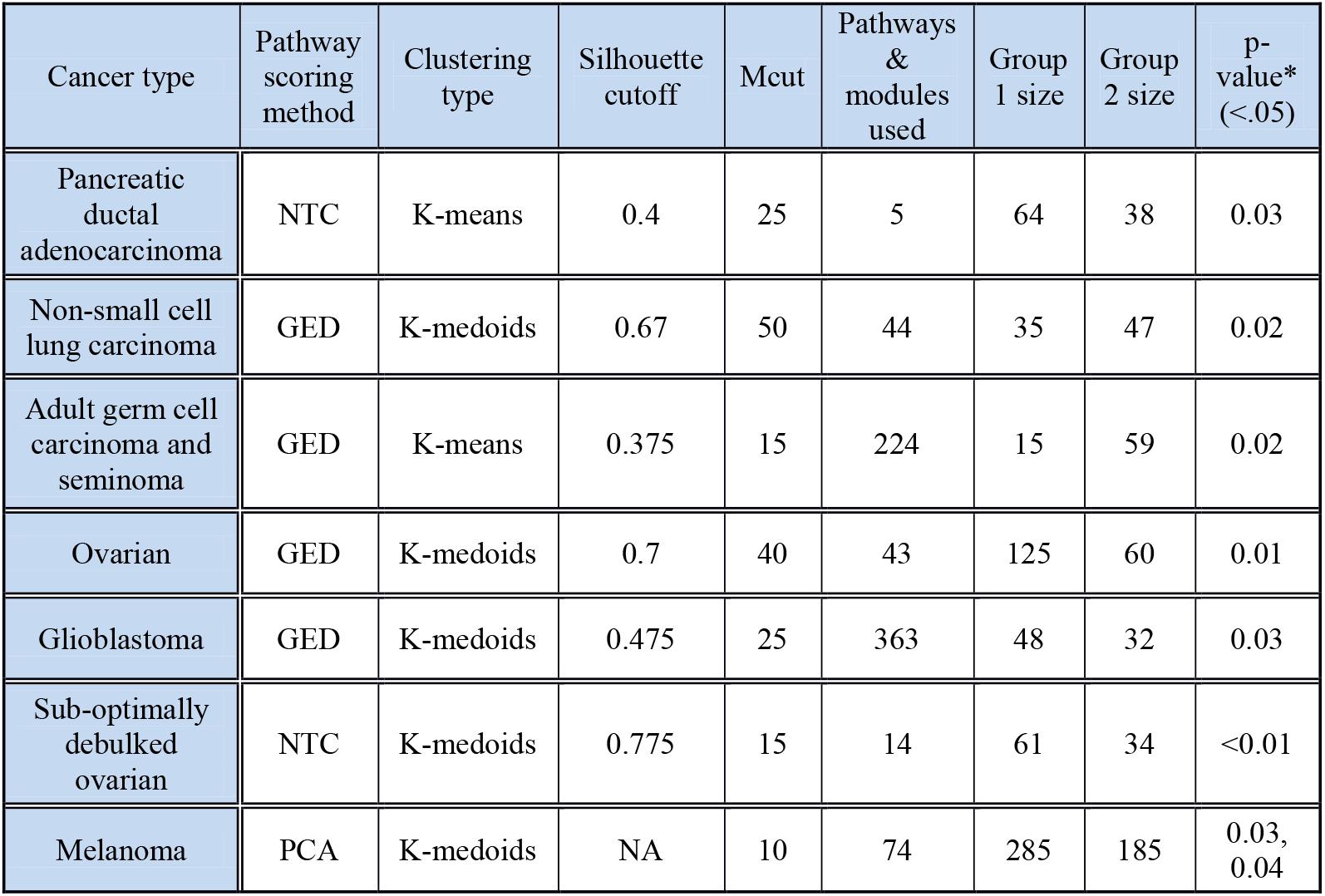
PICS parameter settings and results for individual disease site classifications. Silhouette cuto is for pathway selection and Mcut is for selection of genes in pathways based on their membership in other pathways. P-values are for the logrank statistic for Kaplan-Meier curve separation, and for the melanoma dataset the two values are for the training and the test subsets.

| Cancer type | Pathway scoring method | Clustering type | Silhouette cutoff | Mcut | Pathways & modules used | Group 1 size | Group 2 size | p-value* (<.05) |
| --- | --- | --- | --- | --- | --- | --- | --- | --- |
| Pancreatic ductal adenocarcinoma | NTC | K-means | 0.4 | 25 | 5 | 64 | 38 | 0.03 |
| Non-small cell lung carcinoma | GED | K-medoids | 0.67 | 50 | 44 | 35 | 47 | 0.02 |
| Adult germ cell carcinoma and seminoma | GED | K-means | 0.375 | 15 | 224 | 15 | 59 | 0.02 |
| Ovarian | GED | K-medoids | 0.7 | 40 | 43 | 125 | 60 | 0.01 |
| Glioblastoma | GED | K-medoids | 0.475 | 25 | 363 | 48 | 32 | 0.03 |
| Sub-optimally debulked ovarian | NTC | K-medoids | 0.775 | 15 | 14 | 61 | 34 | <0.01 |
| Melanoma | PCA | K-medoids | NA | 10 | 74 | 285 | 185 | 0.03, 0.04 |

For the lung case, hierarchical clustering of the final pathway matrix yields three distinct groups: the normal samples and groups 1 and groups 2, see Figure 3. This figure, and similar ones in the Supplementary Information, demonstrates that there is greater heterogeneity across tumor samples than across the non-cancerous samples of the same tissue origin. The same groups 1 and 2 were also uncovered via k-medoids clustering into two sets of just the cancer samples, see Figure 4. The clustergram colors (Figure 3) are rescaled automatically for improved visualization of the groups whereas in Figure 4 there is no scaling–the values are directly from the GED pathway scoring. Signaling pathways are the predominant type of pathway used in the lung cancer classification. The general trend, best observed in Figure 4, is that most pathways have both up-regulation of the gene expression in the cancer groups (the bottom right red section) and down-regulation of genes in those same pathways (the upper right blue section). The heatmap in Figure 4 shows a prominent center white strip. The color values in this heatmap correspond to the actual GED values of the pathway scoring (they are not rescaled) and thus we see that for these center strip pathways, the up-regulation color values are close to zero, meaning those pathways, for the cancer samples, are almost exclusively down regulated. These down-regulated pathways for the cancer samples are three endocrine system pathways, two organismal pathways, and two environmental information processing pathways. We also see from both color maps that Group 2, which has less favorable survival statistics, is more distant from the normal gene levels than is Group 1.

**Figure 3:**
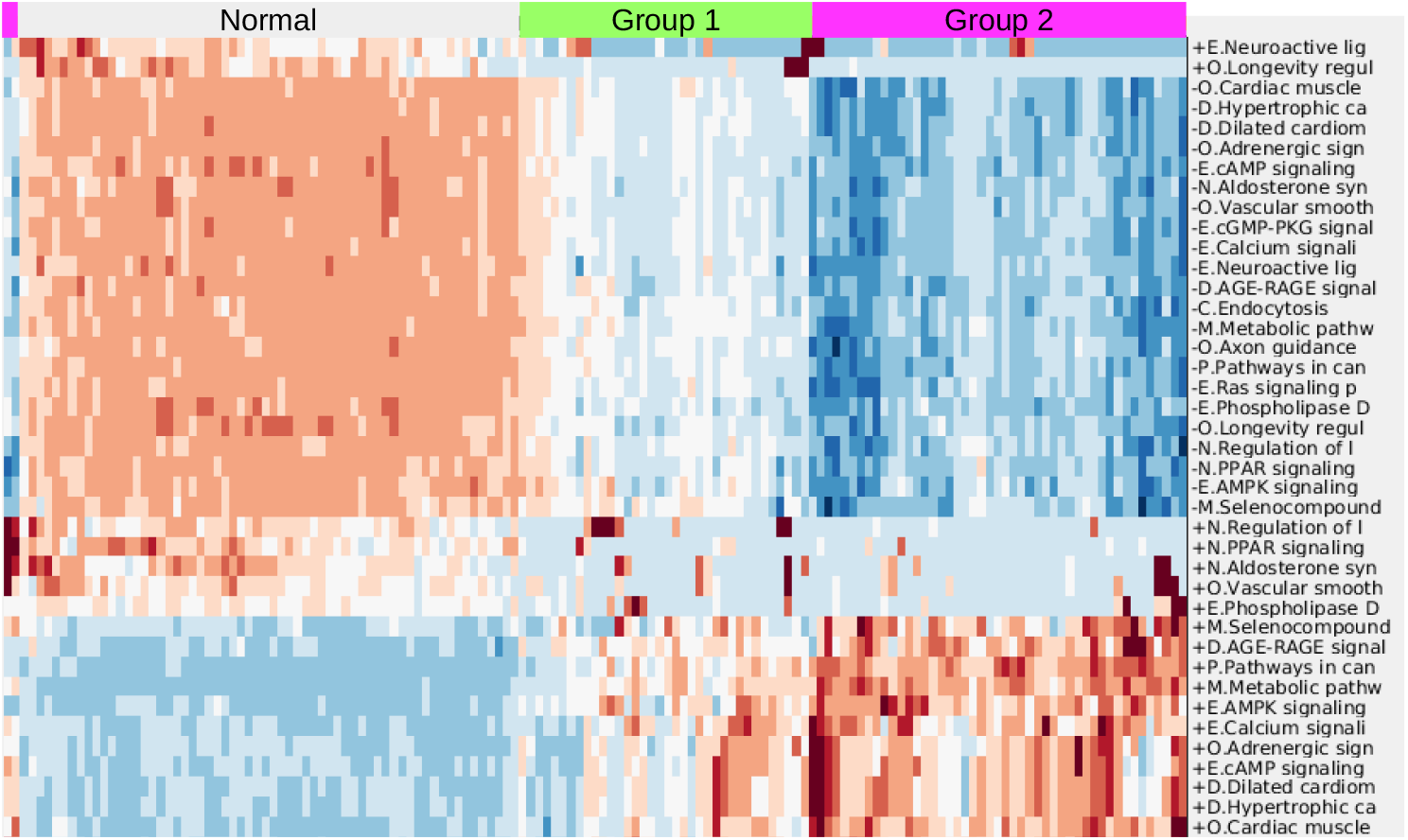
Clustergram for lung case including normal tissues. Colors are automatically scaled for optimal visual distinction, thus the color bar is omitted. For the cancer samples (group 1 and group 2) red indicates increased expression levels of genes in the pathways and blue indicates decreased expression levels. The first letter of each pathway is an abbreviation for the KEGG pathway grouping: E = environmental information processing, O = other organismal systems, D = diseases other than cancer, M = metabolic, C = cellular processes, P = pathways in cancer, and N = endocrine. “+” stands for increased expression of the pathway genes in the cancer samples and “-” stands for decreased expression.

**Figure 4:**
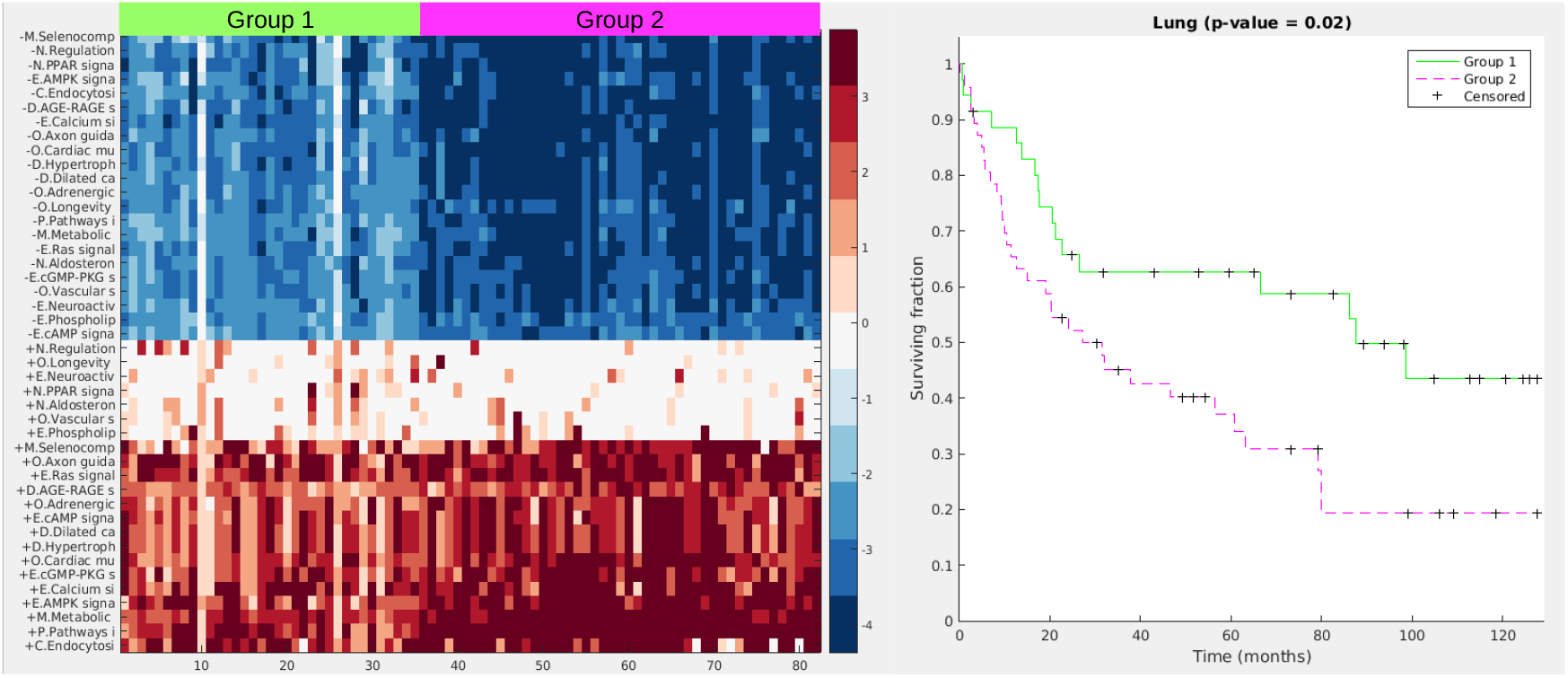
Heatmap and Kaplan-Meier curves (p-value=0.02) for the lung case. Color values are from the pathway scoring by the GED method. Values farther away from 0 mean the pathway gene expression levels are farther away from the gene expression levels of the normal samples.

Figure 5 displays a hybrid heatmap for the pancreas dataset. PICS reduced the 301 pathways and 187 modules from KEGG to a single metabolic pathway (arginine biosynthesis), two cancer pathways (thyroid and endometrial) and two modules (polyamine biosynthesis and urea cycle-1). The pathway signature produced by PICS clustered the patient cohort into three distinct groupings: normal tissue (not shown in heatmap), Group 1, and Group 2. Note that cancer pathways curated in KEGG, such as the thyroid cancer and the endometrial cancer, are in fact relatively general cancer pathways, containing the signaling sub-pathways commonly implicated in cancer, including MAPK, RAS, PI3K-AKT, Wnt, p53, and ErbB. The heatmap shows that Group 2, which has worse survivability, is more dysregulated, as judged by NTC score, than Group 1. Since with the NTC method it is not possible to directly assess increased or decreased gene expression levels for pathways in the heatmap figure, Figure 6 displays the gene expression levels across the groups for one of the pathways used in the pancreas clustering, the thyroid cancer pathway. Here we observe that most genes in this pathway are on average similar across the three groups, with a tendency towards more variability in Group 2, but some genes, such as MAPK1, PAX8, and CCDC6, are up-regulated.

**Figure 5:**
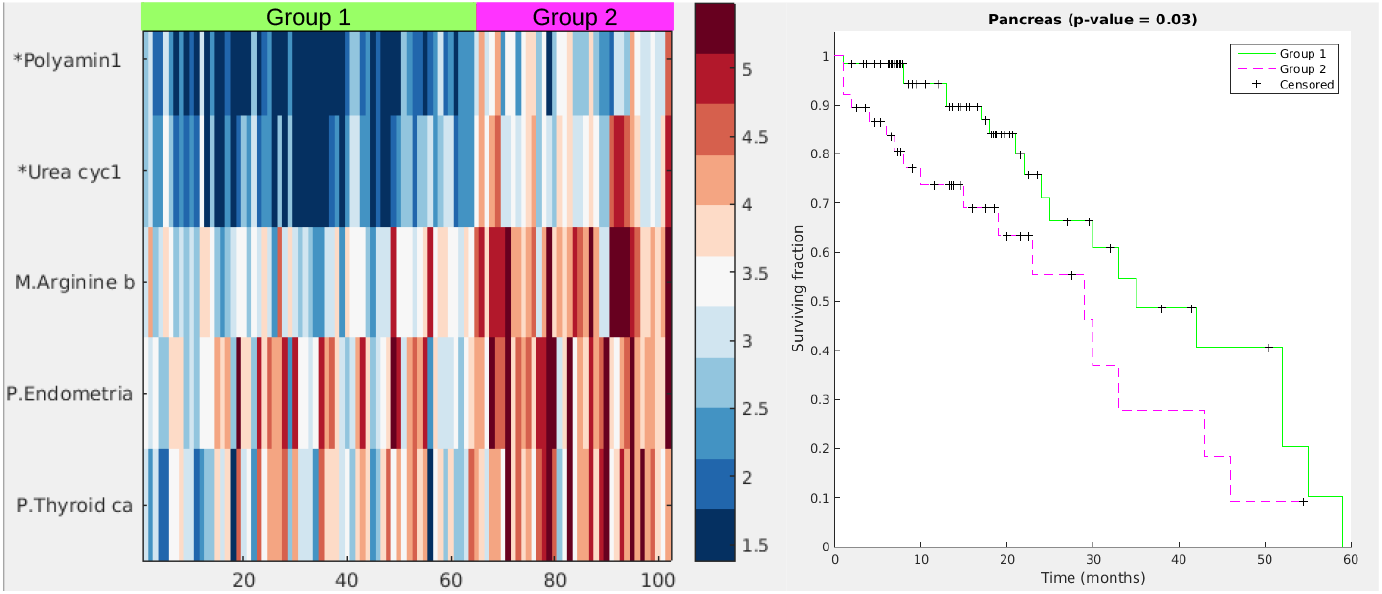
Heatmap and Kaplan-Meier curves (p-value=0.03) for the pancreas case. Color values are from the NTC pathway scoring method, which yields higher values for the farther away a pathway’s gene expression levels are from the gene levels of the normal samples. Group 2 is farther away in general than Group 1 and also has worse survivability, similar to the lung case.

**Figure 6:**
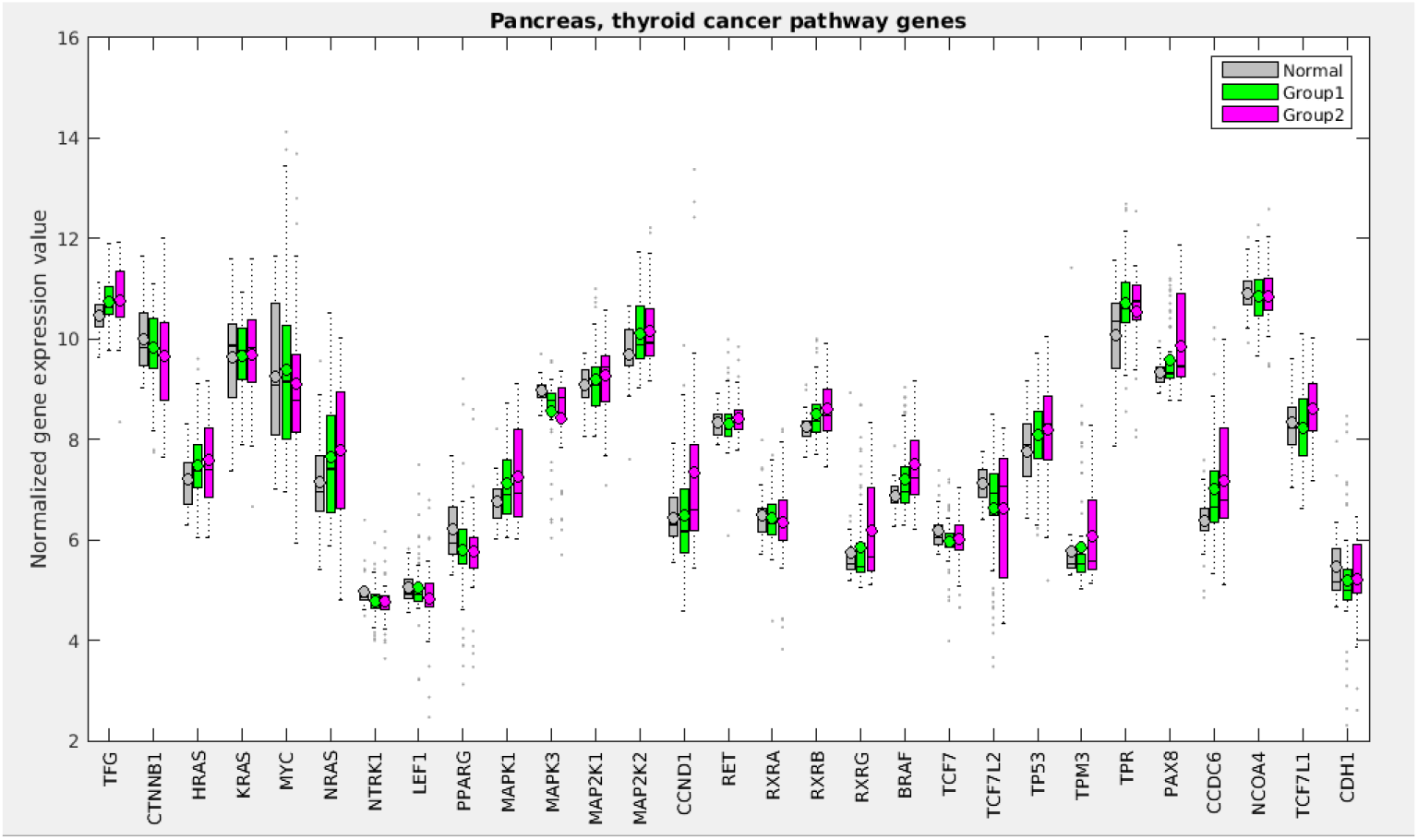
Gene expression levels for the pancreas cohort for the genes in the thyroid cancer pathway.

See the Supplementary Information for the heatmaps and Kaplan-Meier curves for the other individual disease sites.

### 3.3 Individual cancer site clustering without normal samples

Figure 7 shows the KM curve separation for this run for the training and the test sets.

**Figure 7:**
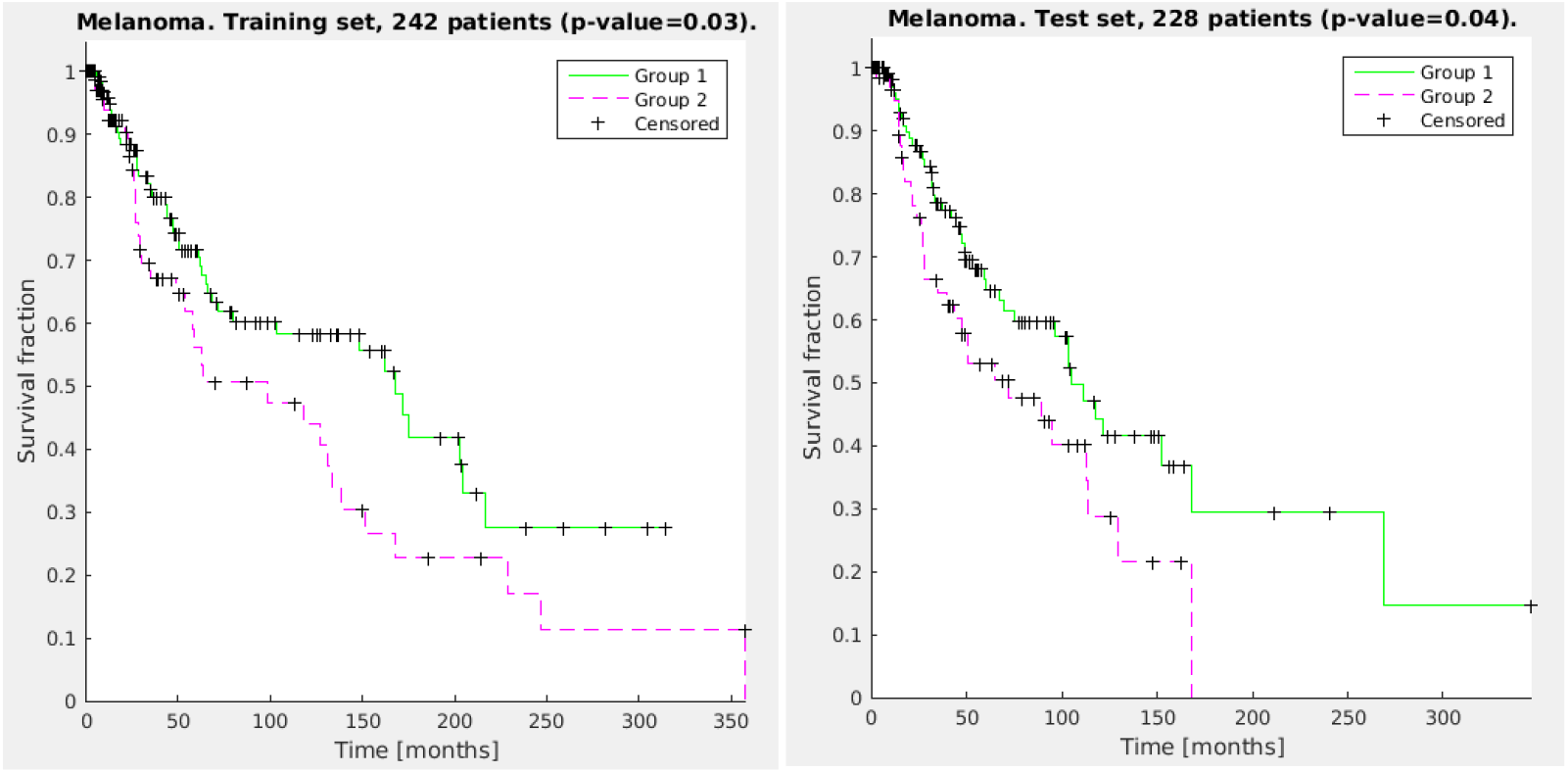
Kaplan-Meier curves for a representative run from the optimal parameter set found for the melanoma dataset. Both the training set and the test set show statistically significant separation between the two clustered groups.

Figure 8 displays the resulting pathway matrix for the melanoma run as a heatmap, and contains the entire dataset (training and test). The most prominent pathway groupings are the thick red-blue band towards the top and the thinner blue-red band at the bottom. Investigation into the pathways involved in these bands (which are not shown due to the large number of pathways used, but see the Supplementary Information for the list of pathways in the same order as the heatmap rows) reveals that these are immune system pathways, which aligns with the general opinion of the relevance of the immune system in melanoma [26]. Group 1, which has generally higher levels of active immune system genes–as demonstrated in Figure 9 which shows the gene expression comparison for the T-cell receptor signaling pathway–also shows better survival characteristics.

**Figure 8:**
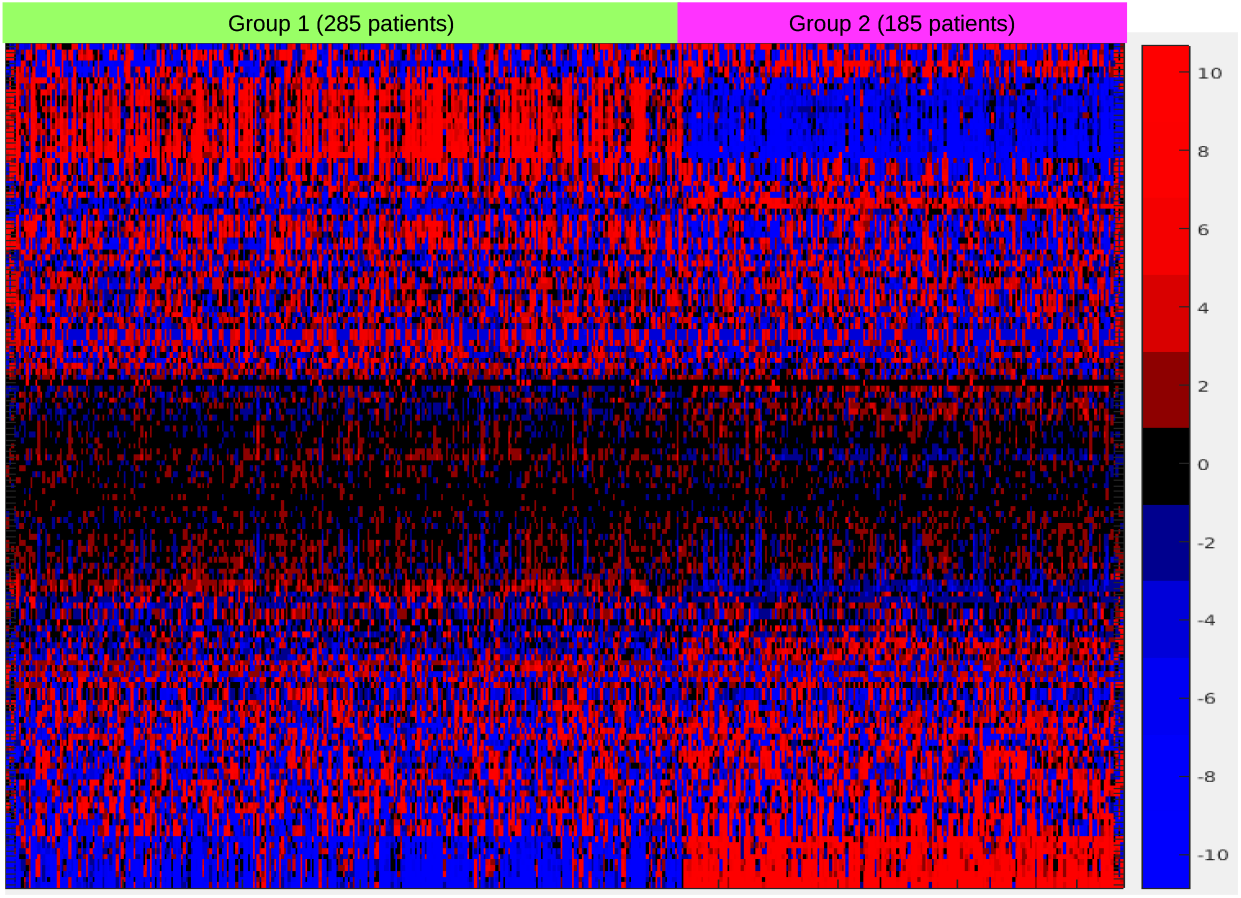
Hierarchical clustering for the pathways and k-medoids clustering for the combined training and testing samples. Pathway labels have been suppressed since 148 pathways and modules were used. The prominent red-blue band towards the top and blue-red band at the bottom are dominated by immune system-related pathways, and investigation into the gene expression levels from those pathways reveals heightened immune system expression for group 1, which showed higher overall median survival. The color scale values are the values from the PCA and are thus not directly interpretable.

## 4 Discussion and conclusions

The heterogeneity across cancers, even within the same type of cancer, has been discussed much in the literature over the past decade [27]. However, for the most part it has not been known how to use this information to better inform treatment decisions for patients [28]. In this report we have introduced a method called PICS that can classify cancer patients based on their gene expression levels by compressing this information from the gene level to the pathway level. We view this data regularization/dimension reduction technique and the accompanying classification system as the first step in building a computational tool to aid with patient treatment planning decision making. PICS is advantageous due to its speed (a PICS run takes minutes compared to the related method Pathifier [10] which takes hours), its interpretability, and its flexibility to handle a variety of types of datasets, as we have demonstrated. Although our main focus is broader and more statistical in nature, we welcome deeper biological analysis on the multitude of discovered pathway-cancer corollaries.

**Figure 9:**
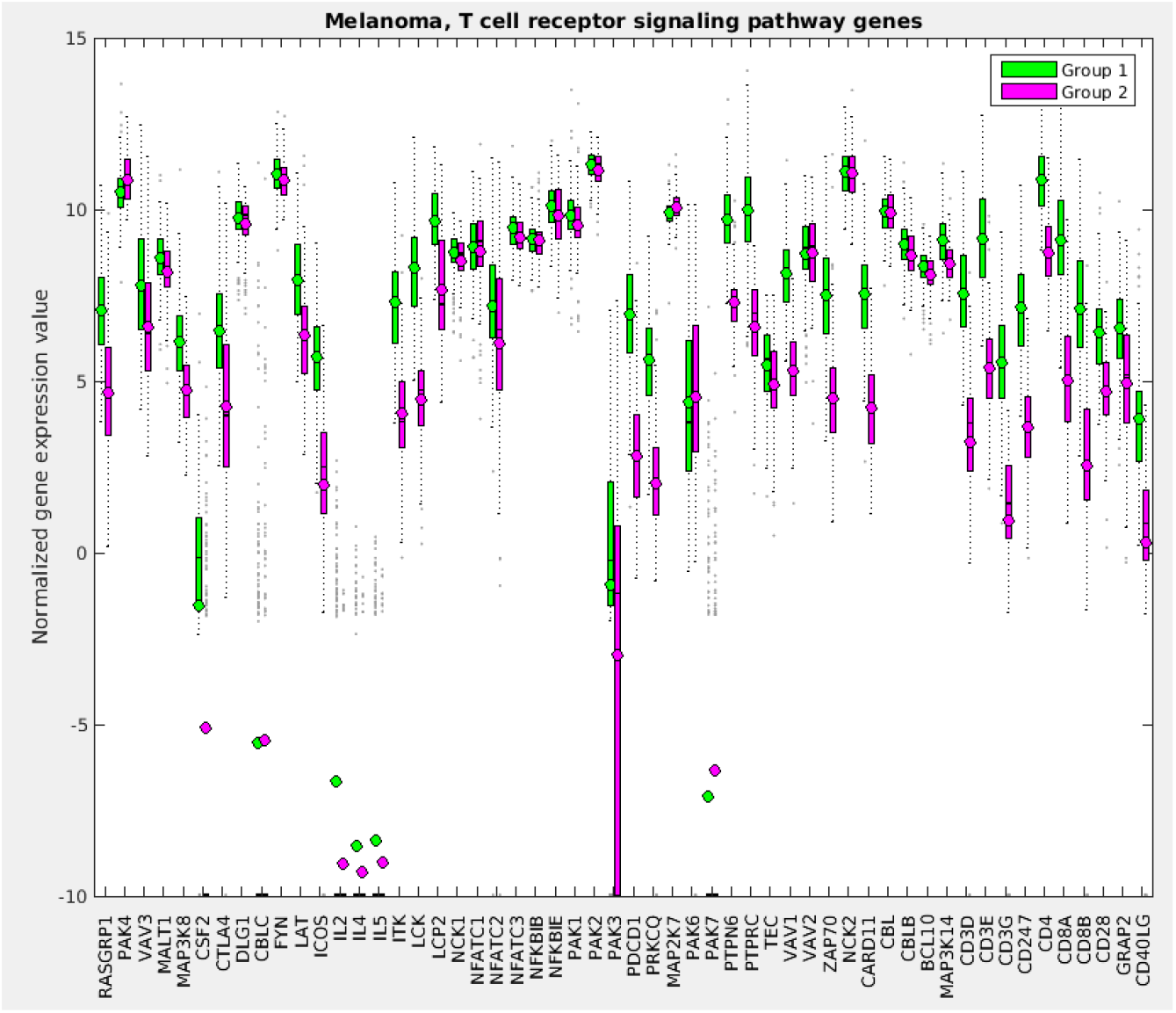
A sample gene expression value plot for the melanoma training set of the T-cell receptor signaling pathway showing that in general group 1, the higher survivability class, has elevated expression levels for the genes involved in that pathway.

The ultimate vision is a system that takes into account a plethora of patient-level information (tumor site, stage, size, tumor and germline genomics data, patient age, etc.) and predicts what drugs and drug levels (including radiation) will likely be most beneficial and least toxic to the patient. With this goal in mind, we began with a method that classifies patients based on only their gene expression data, as measured by microarrays (RNAseq data would also be immediately possible to use). We have shown across a wide range of cancer types that classification based on biological pathway distillation of gene expression data allows for grouping patients, via clustering algorithms, into distinct survivability classes. The next steps are to obtain datasets containing treatment information as well as genomic and clinical variables, either by mining currently existing sources, by performing clinical trials, or through institutional data gathering, in order to build a machine learning system for outcome prediction. We hypothesize that compressing the genetic information into pathway level scores will be useful in this broader context.

While we have explored several variants of the PICS algorithm for classifying the various cancer types studied, there is still room for refinement in the technique. For example, we relied solely on KEGG curated pathways. It remains to be investigated if other curated pathway databases might be better suited to the task. One shortcoming in KEGG is that the redundancy across pathways since many pathways are built up of other smaller pathways. The MSigDB [29] database, which consolidates many pathway resources into a single repository, will likely be a useful resource. It also remains to be seen if using the network connectivity of the pathways rather than just the list of genes the pathways contain will be a useful adjustment to the algorithm. We hope that by building a method that allows for pan-cancer analysis, we can extend learning across cancer types and make cancer research less of an isolated, site-specific endeavor. If common pathway signatures across various cancer types end up being predictive of the success of drugs, the goals of biological knowledge building and improved patient care will both have been achieved.

There are pros and cons associated with each of the pathway scoring methods introduced. The PCA method is the simplest method, and is applicable to datasets that do not have normal tissues, but it was outperformed in the datasets that did contain normal tissues. This is likely because both the NTC and GED methods score pathways based on how different they are from the normal pathways, and this information, not surprisingly, is useful for classifying cancer patients. Our results echo the trend that has been observed before that the more dysregulation there is across pathways, the more malignant is the cancer, e.g. [10, 30]. It may prove useful to combine different scoring methods in the same analysis, for example some pathways might better be scored by GED and some better by NTC, but we have not investigated that as of yet. We also used the global parameter Mcut to vary how many genes are used to score each of the pathways, but the scoring of each pathway should likely be done in a more pathway-dependent manner. Nevertheless, the results show good classification even without detailed modeling of pathway activity. We have intentionally tried to find the right balance between statistical modeling and biology in this effort, in order to not be pulled too far in either direction.

The most important genomic addition to PICS will be the ability to include more molecular information, including post-transcriptional and post-translational data, into the pathway scoring. We chose pathways as our fundamental unit for classification based on the idea that they represent the right level of biological detail for the statistical prediction problem, and pathway activity can be informed by a large variety of genomic assays. Specifically, we expect gene mutations, chromosomal rearrangements and fusions, copy number variations, methylation, and proteomics to all yield useful information regarding pathway activity. Our current pathway scoring and clustering approach will also benefit from insights into biological redundancies and more complex modeling of cellular and organismal functioning. For example, the use of logical statements like AND, OR, IF, etc. in modeling cellular systems will likely aid in classification, but this makes the search space for optimal classifiers combinatorial and therefore much harder. Informing these additions with known experimental biology will be necessary.

Although we stressed classifying cancers in this report, we also plan to apply PICS to classify normal tissues and the relevant organ systems of patients in order to build a system to predict toxicities and side effects as well, which is equally important for optimal cancer management. In the last five years, 70 cancer drugs were approved by the FDA. At this rate of oncology drug approval, sophisticated clinical decision tools are increasingly needed to make the best therapeutic choices.

## References

[1] David W Scott, Fong Chun Chan, Fangxin Hong, Sanja Rogic, King L Tan, Barbara Meissner, Susana Ben-Neriah, Merrill Boyle, Robert Kridel, Adele Telenius, et al. Gene expression– based model using formalin-fixed paraffin-embedded biopsies predicts overall survival in advanced-stage classical hodgkin lymphoma. Journal of Clinical Oncology, 31(6):692–700, 2013.

[2] Sandy D Der, Jenna Sykes, Melania Pintilie, Chang-Qi Zhu, Dan Strumpf, Ni Liu, Igor Jurisica, Frances A Shepherd, and Ming-Sound Tsao. Validation of a histology-independent prognostic gene signature for early-stage, non–small-cell lung cancer including stage ia patients. Journal of Thoracic Oncology, 9(1):59–64, 2014.

[3] Anastasia Chalkidou, Michael J ODoherty, and Paul K Marsden. False discovery rates in PET and CT studies with texture features: a systematic review. PloS one, 10(5):e0124165, 2015.

[4] Minoru Kanehisa, Yoko Sato, Masayuki Kawashima, Miho Furumichi, and Mao Tanabe. KEGG as a reference resource for gene and protein annotation. Nucleic acids research, 44(D1):D457–D462, 2016.

[5] Darryl Nishimura. Biocarta. Biotech Software & Internet Report: The Computer Software Journal for Scient, 2(3):117–120, 2001.

[6] G Joshi-Tope, Marc Gillespie, Imre Vastrik, Peter D’Eustachio, Esther Schmidt, Bernard de Bono, Bijay Jassal, GR Gopinath, GR Wu, Lisa Matthews, et al. Reactome: a knowledgebase of biological pathways. Nucleic acids research, 33(supp 1):D428–D432, 2005.

[7] Alexander R Pico, Thomas Kelder, Martijn Pvan Iersel, Kristina Hanspers, Bruce R Conklin, and Chris Evelo. Wikipathways: pathway editing for the people. PLoS biology, 6(7), 2008.

[8] C. Vaske, S. Benz, J. Sanborn, D. Earl, C. Szeto, J. Zhu, D. Haussler, and J. Stuart. Inference of patient-specific pathway activities from multi-dimensional cancer genomics data using PARADIGM. Bioinformatics, 26(12):i237–i245, 2010.

[9] Maysson Ibrahim, Sabah Jassim, Michael A Cawthorne, and Kenneth Langlands. A matlab tool for pathway enrichment using a topology-based pathway regulation score. BMC bioinformatics, 15(1):358, 2014.

[10] Y. Drier, M. Sheffer, and E. Domany. Pathway-based personalized analysis of cancer. Proceedings of the National Academy of Sciences, 110(16):6388–6393, 2013.

[11] A. Tarca, S. Draghici, P. Khatri, S. Hassan, P. Mittal, J. Kim, C. Kim, J. Kusanovic, and R. Romero. A novel signaling pathway impact analysis. Bioinformatics, 25(1):75–82, 2009.

[12] A. Margolin, I. Nemenman, K. Basso, C. Wiggins, G. Stolovitzky, R. Favera, and A. Califano. ARACNE: an algorithm for the reconstruction of gene regulatory networks in a mammalian cellular context. BMC bioinformatics, 7(supp 1):S7, 2006.

[13] N. Duarte, S. Becker, N. Jamshidi, I. Thiele, M. Mo, T. Vo, R. Srivas, and B. Palsson. Global reconstruction of the human metabolic network based on genomic and bibliomic data. Proceedings of the National Academy of Sciences, 104(6):1777–1782, 2007.

[14] Shinuk Kim, Mark Kon, and Charles DeLisi. Pathway-based classification of cancer subtypes. Biology Direct, 7(1):21, 2012.

[15] L. Van der Maaten and G. Hinton. Visualizing data using t-SNE. Journal of Machine Learning Research, 9(2579-2605):85, 2008.

[16] E. Levina and P. Bickel. Maximum likelihood estimation of intrinsic dimension. In Advances in neural information processing systems, pages 777–784, 2004.

[17] Leonard Kaufman and Peter J Rousseeuw. Finding groups in data: an introduction to cluster analysis, volume 344. John Wiley & Sons, 2009.

[18] Asoke K Nandi, Rui Fa, and Basel Abu-Jamous. Integrative Cluster Analysis in Bioinformatics. John Wiley & Sons, 2015.

[19] A. Gentles, A. Newman, C. Liu, S. Bratman, W. Feng, D. Kim, V. Nair, Y. Xu, A. Khuong, C. Hoang, and et al. The prognostic landscape of genes and infiltrating immune cells across human cancers. Nature medicine, 21(8):938–945, 2015.

[20] Jeran K Stratford, David J Bentrem, Judy M Anderson, Cheng Fan, Keith A Volmar, JS Mar-ron, Elizabeth D Routh, Laura S Caskey, Jonathan C Samuel, Channing J Der, et al. A six-gene signature predicts survival of patients with localized pancreatic ductal adenocarcinoma. PLoS Med, 7(7):e1000307, 2010.

[21] Panagiotis A Konstantinopoulos, Dimitrios Spentzos, Beth Y Karlan, Toshiyasu Taniguchi, Elena Fountzilas, Nancy Francoeur, Douglas A Levine, and Stephen A Cannistra. Gene expression profile of brcaness that correlates with responsiveness to chemotherapy and with outcome in patients with epithelial ovarian cancer. Journal of Clinical Oncology, 28(22):3555–3561, 2010.

[22] James E Korkola, Jane Houldsworth, Rajendrakumar SV Chadalavada, Adam B Olshen, Debbie Dobrzynski, Victor E Reuter, George J Bosl, and RSK Chaganti. Down-regulation of stem cell genes, including those in a 200-kb gene cluster at 12p13. 31, is associated with in vivo differentiation of human male germ cell tumors. Cancer research, 66(2):820–827, 2006.

[23] Tomas Bonome, Douglas A Levine, Joanna Shih, Mike Randonovich, Cindy A Pise-Masison, Faina Bogomolniy, Laurent Ozbun, John Brady, J Carl Barrett, Jeff Boyd, et al. A gene signature predicting for survival in suboptimally debulked patients with ovarian cancer. Cancer research, 68(13):5478–5486, 2008.

[24] Yohan Lee, Adrienne C Scheck, Timothy F Cloughesy, Albert Lai, Jun Dong, Haumith K Farooqi, Linda M Liau, Steve Horvath, Paul S Mischel, and Stanley F Nelson. Gene expression analysis of glioblastomas identifies the major molecular basis for the prognostic benefit of younger age. BMC medical genomics, 1(1):52, 2008.

[25] Tomas Bonome, Douglas A Levine, Joanna Shih, Mike Randonovich, Cindy A Pise-Masison, Faina Bogomolniy, Laurent Ozbun, John Brady, J Carl Barrett, Jeff Boyd, et al. A gene signature predicting for survival in suboptimally debulked patients with ovarian cancer. Cancer research, 68(13):5478–5486, 2008.

[26] Cancer Genome Atlas Network et al. Genomic classification of cutaneous melanoma. Cell, 161(7):1681–1696, 2015.

[27] R Fisher, L Pusztai, and C Swanton. Cancer heterogeneity: implications for targeted therapeutics. British journal of cancer, 108(3):479–485, 2013.

[28] Teri A Manolio. Bringing genome-wide association findings into clinical use. Nature Reviews Genetics, 14(8):549–558, 2013.

[29] Arthur Liberzon, Aravind Subramanian, Reid Pinchback, Helga Thorvaldsd´ottir, Pablo Tamayo, and Jill P Mesirov. Molecular signatures database (MSigDB) 3.0. Bioinformatics, 27(12):1739–1740, 2011.

[30] S Rondeau, S Vacher, L De Koning, A Briaux, A Schnitzler, W Chemlali, C Callens, R Lidereau, and I Bièche. ATM has a major role in the double-strand break repair pathway dysregulation in sporadic breast carcinomas and is an independent prognostic marker at both mrna and protein levels. British journal of cancer, 112(6):1059–1066, 2015.

